# A systematic comparison of predictive models on the retina

**DOI:** 10.1101/2024.03.06.583740

**Authors:** Michaela Vystrčilová, Shashwat Sridhar, Max F. Burg, Mohammad H. Khani, Dimokratis Karamanlis, Helene M. Schreyer, Varsha Ramakrishna, Steffen Krüppel, Sören J. Zapp, Matthias Mietsch, Tim Gollisch, Alexander S. Ecker

**Author notes:** **Data availability:** All data and code will be made available upon publication. **Competing interests:** The author declare no competing interests.

## Abstract

Understanding the nonlinear encoding mechanisms of retinal ganglion cells (RGCs) in response to various visual stimuli presents a central challenge in neuroscience, driving the development of increasingly complex predictive models. Here, we systematically evaluate linear-nonlinear (LN) models – applying various regularization techniques – and convolutional neural networks (CNNs) of increasing depth, to predict RGC responses to white noise and natural movies. Our analysis includes publicly available datasets from marmoset and salamander retinas.

We demonstrate that LN models, when equipped with appropriate inductive biases, can achieve robust predictive performance on neural responses to both white noise and natural movie stimuli. The optimal inductive biases vary substantially across datasets and stimulus types, indicating that the LN model’s performance is susceptible to these choices. This warrants care when using LN models as baselines: their performance is not fixed, and inappropriate design choices can lead to “unfair” comparisons.

However, even in the optimal inductive bias scenario, CNNs consistently outperform LN models across conditions, confirming the advantage derived from their nonlinear representation capacity. Investigating cross-stimulus generalization, we observe that models trained on white noise generalize better to natural movies than vice versa. Notably, LN models exhibit a smaller performance gap between in-domain and cross-domain predictions compared to CNNs, suggesting that the nonlinear processing captured by CNNs is more stimulus-specific.

Overall, this study provides valuable benchmarks and methodological insights for neuroscientists designing predictive models of retinal encoding.

## Introduction

Predictive models are fundamental tools in systems neuroscience because they transform qualitative ideas about neural coding into quantitative hypotheses whose success or failure can be directly measured against data (Rieke et al., 1999; Dayan and Abbott, 2005). By estimating the mapping from sensory input to spiking output, such models enable the testing of competing theories of computation, the design of closed-loop experiments, and the development of interfaces between brains and machines.

Classical linear-nonlinear (LN) models are a well-established model for linearly characterizing RGC responses (Marmarelis and Naka, 1972; Korenberg and Hunter, 1986; Sakai et al., 1988; Chichilnisky, 2001; Pillow et al., 2008). However, they cannot capture their nonlinear computations such as contrast adaptation, motion anticipation, and omitted stimulus response (Schwartz et al., 2007, 2012; Kastner and Baccus, 2014; Karamanlis and Gollisch, 2021; Maheswaranathan et al., 2023). These limitations have been addressed through augmenting such models with nonlinear, yet interpretable features that capture RGC responses more accurately (Baccus et al., 2008; McFarland et al., 2013; Liu et al., 2017; Shi et al., 2019; Sridhar et al., 2025).

More recently, deep learning models, particularly convolutional neural networks (CNNs), have emerged as a tool for predicting RGC responses, outperforming LN models (Batty et al., 2016; McIntosh et al., 2016). Besides improving prediction accuracy, CNNs also facilitated new insights into retinal cell types (Höfling et al., 2024), and have helped develop mechanistic theories of how nonlinear computations in the retina are implemented (Tanaka et al., 2019; Maheswaranathan et al., 2023). While LN models have been challenged for their often poor response prediction to natural stimuli (Heitman et al., 2016; McIntosh et al., 2016), CNNs can predict responses to natural stimuli well (Batty et al., 2016; McIntosh et al., 2016; Höfling et al., 2024).

Open comparison has driven model improvement in machine learning. Computational neuroscience has begun to adopt the same strategy. One of the earliest examples was the Berkeley Neural Prediction Challenge^1^, which released public training data for neurons recorded in the primary visual cortex, the primary auditory cortex, and the songbird field L. Since then, several initiatives have appeared – Sensorium (Willeke et al., 2022; Turishcheva et al., 2023) for the mouse visual cortex, Brain-Score (Schrimpf et al., 2020) for the primate ventral stream, Neural Latents ‘21 (Pei et al., 2021) for sensory, motor, and cognitive monkey cortical areas, and the Algonauts series (Cichy et al., 2019) for human fMRI and MEG data. However, a quantitative benchmark of models predicting single-neuron responses of RGCs is still lacking.

The generalization of predictive models across stimuli has also been the subject of recent studies. In V1, it has been shown that models trained on natural movie stimuli generalize well to white noise (Sinz et al., 2018) but not the other way around (David et al., 2004; Sinz et al., 2018). In the context of the retina, poor generalization from white noise to natural movies has been reported (Heitman et al., 2016; McIntosh et al., 2016). It has been attributed to white noise lacking essential statistical features of natural scenes and therefore failing to capture important nonlinear computations (Maheswaranathan et al., 2023). Vystrčilová et al. (2025) showed the opposite – that LN models generalize better from white noise to natural movies than the other way around. However, neither provided a systematic study of generalization abilities across stimuli and models of varying complexities in the retina.

In this paper, we address these gaps by comparing models of varying complexities across white noise and natural stimuli. We evaluate their suitability for modeling RGC responses and their generalization across stimuli, serving as a reference point, providing strong baselines for future research. Specifically, we first train and evaluate LN and CNN models on both white noise and natural movie stimuli. We show that LN models can perform surprisingly well on both stimulus types when equipped with appropriate regularization and carefully chosen inductive biases. However, the effectiveness of these models depends strongly on the specific biases used, which differ across datasets and stimulus conditions. This suggests that LN models are not fixed baselines and that their performance can vary considerably depending on design choices. Nonlinear multi-layer CNNs, nonetheless, show better predictive capabilities across species and stimuli.

Secondly, we investigate whether training on one of the stimulus types is sufficient for the model to generalize to the other, and if so, at what model complexity. We find that in neither case do the models reach in-domain performance: the model trained and evaluated on the same stimulus type always outperforms the one trained on one stimulus type and evaluated on the other. This suggests that the underlying retinal processing adapts to the statistics of the stimulus. We observe that generalization works better from models trained on white noise to natural movies than the other way around, consistent with the results of Vystrčilová et al. (2025). Interestingly, the LN models approach their in-domain performance more closely than the nonlinear models, suggesting that it is predominantly the nonlinear computation of the retina that adjusts between the processing of the two stimuli.

## Results

We analyzed the activity of marmoset (*Callithrix jacchus*) and salamander (*Ambystoma mexicanum*) retinal ganglion cells (RGCs) in response to white noise (WN) and natural movie (NM) visual stimuli recorded using microelectrode arrays (Fig. 1). Our datasets contained the activity of 475 RGCs across four marmoset retinas, which were shown both natural movie and white noise stimuli, and the activity of 186 RGCs across five salamander retinas, which were shown only white noise stimuli. We selected these cells based on the reliability of their responses.

**Figure 1.**
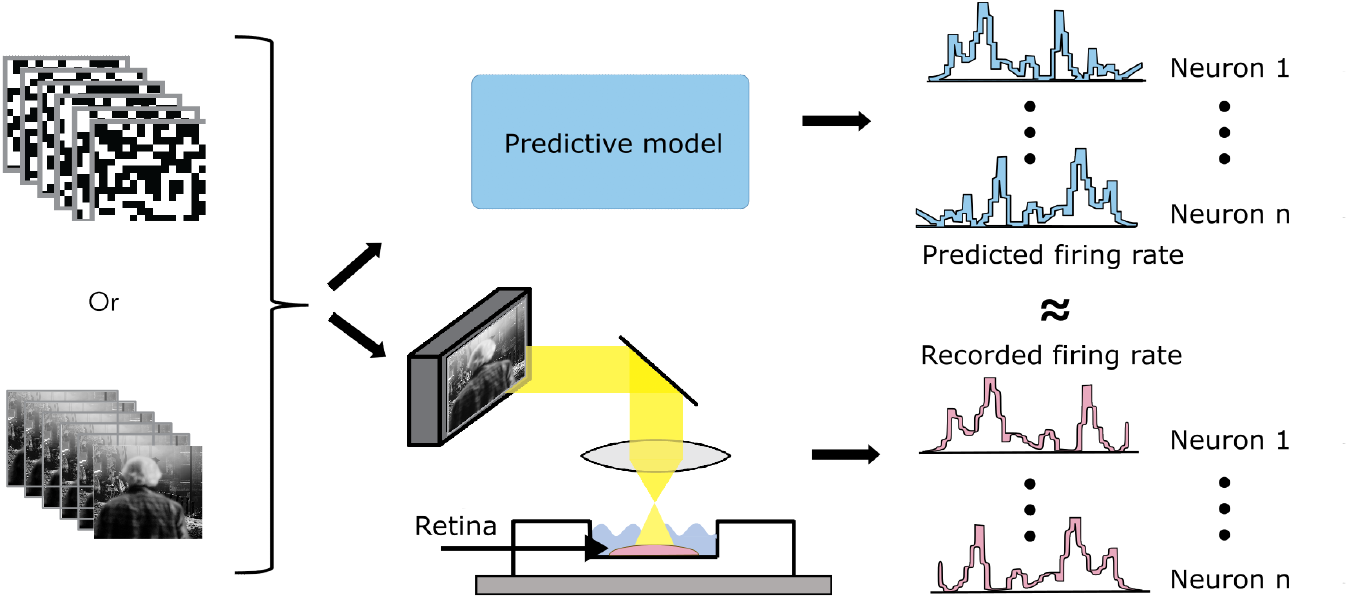
We present white noise or natural movie stimuli to a retina while recording ganglion cell responses using a multielectrode array. We train predictive models (LN and CNN models) to predict the average firing rate of all neurons given the stimulus within a time window preceding the prediction.

The stimuli were split into non-repeating, unique sequences used for training models (“training set”) and a smaller number of repeated segments for evaluating the final model performance (“test set”). A sequence of a non-repeating segment followed by a repeated segment comprised a “trial”. In our data, there were 10 to 248 stimulus trials per retina. The non-repeating segments had 50–300 seconds, the repeating segments 10–60 seconds, depending on the stimulus type and species.

We trained LN and CNN models to predict the neural responses in these datasets. To evaluate the performance of our models, we computed the correlation coefficient between the predicted spike counts and the trial-averaged neuronal response, binned per frame, on the test segments (Equation 3).

### Linear-nonlinear models are useful predictive model benchmarks on white noise as well as natural movie stimuli

LN models are a commonly used approach for modeling RGC responses to white noise. We begin by establishing their predictive performance not only on white noise but also on natural movie stimuli to determine whether they can predict RGC responses to movie stimuli and serve as a baseline for more complex, nonlinear models in our analyses. The LN models consist of a linear spatio-temporal filter capturing the neuron’s linear receptive field, followed by a nonlinearity. To obtain the neurons’ spiking responses, we took the inner product between the stimulus and this linear filter and applied a standard softplus nonlinearity (Fig. 2A).

**Figure 2.**
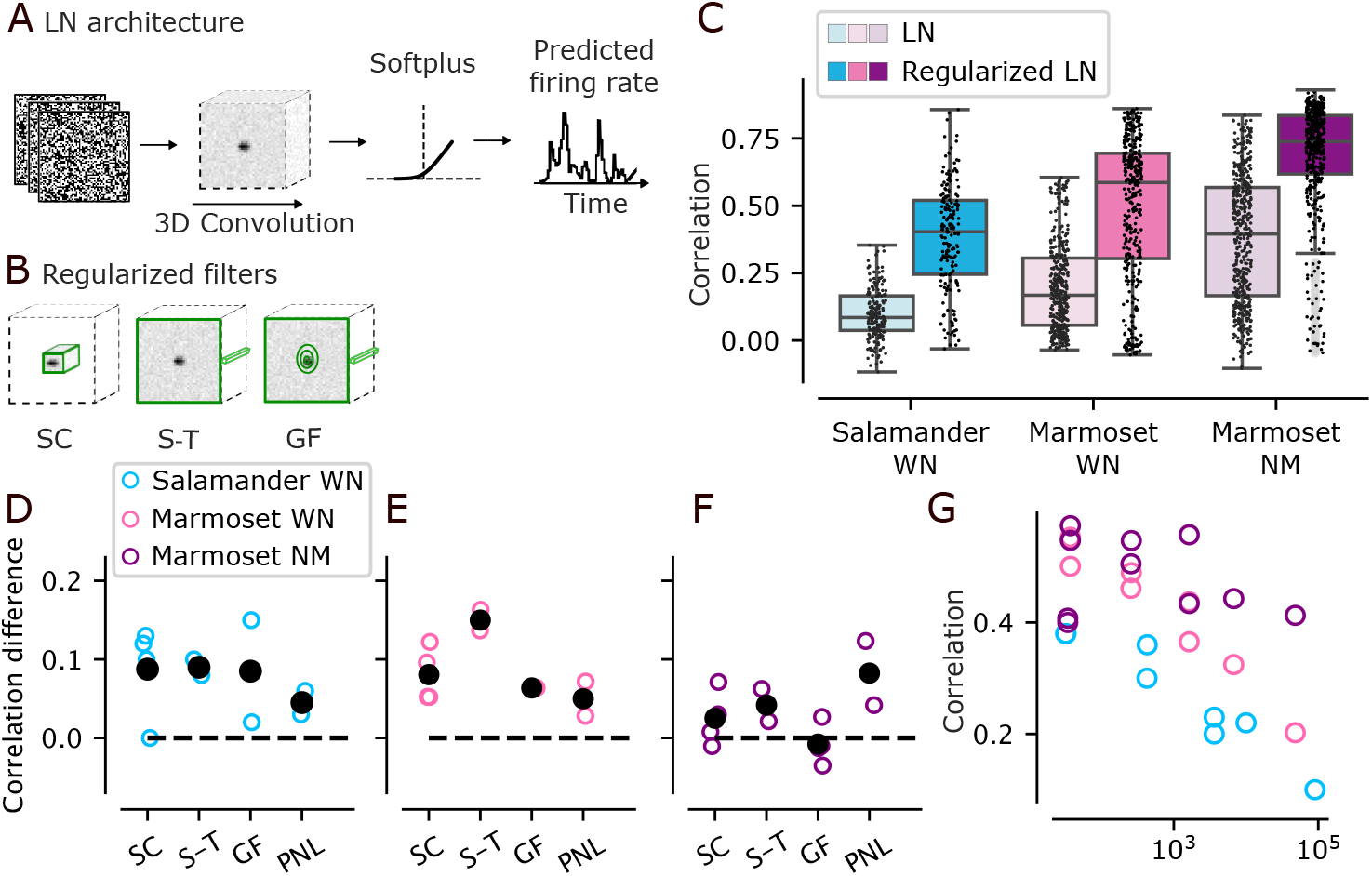
**A**. Basic LN model architecture without any of the regularization methods applied. **B**. Green outlines show how the LN model kernels change based on the regularization methods – *spatial-crop (SC), space-time separation (S-T), Gaussian fitting (GF)*. **C**. Performance comparison between non-regularized LN and one with the best combination of regularizers (inductive biases). **D.–F**. Each colored circle represents the difference that adding the inductive bias on the x-axis makes in performance when keeping all other biases constant. The black dots represent the mean improvement for a given inductive bias across all scenarios. **G**. Each circle is a single model’s performance vs. the number of parameters it has.

A simple way of obtaining linear filters for white noise stimuli is spike-triggered averaging (STA), which we used for the noise stimulus models. For natural movie stimuli, the STA does not recover the correct filter due to the correlations within the stimulus. We, therefore, fit the LN model directly using maximum likelihood estimation by stochastic gradient descent.

A basic, unregularized LN model architecture resulted in a median correlation of 0.09 (IQR 0.05–0.17) (Equation 3) over cells on salamander noise data, 0.18 (IQR 0.1–0.31) of median correlation over cells on marmoset noise data and 0.4 of median (IQR 0.17–0.59) correlation over cells on marmoset movie data. The marmoset natural movie models performed notably better, presumably because a larger fraction of the response variance to natural movies is caused by local brightness fluctuations, which can be captured well by a linear model.

We suspected that the relatively poor performance of this basic LN architecture was caused by overfitting, as it has too many parameters for the given amount of recorded data. To prevent overfitting, we turned to regularization – reducing the number of parameters or adding smoothing priors that restrict the values of the parameters. We used various regularization techniques on the LN models to improve their performance. All of them can be viewed as inductive biases that we impose on our model based on known properties of receptive fields. We examined how strongly different regularization methods which are commonly used when fitting LN models affect the performance, namely: (1) *spatial-crop (SC)*: cropping of the model’s spatial weights to the neurons’ STA-based receptive fields obtained from white noise, (2) *space-time separation (S-T)*: decomposing the spatio-temporal filter into separate spatial and temporal filters, and (3) *Gaussian fitting (GF)*: fitting a two-dimensional Gaussian to the spatial component of the space-time separated filter of the models (Fig. 2B). For noise stimuli, Gaussian fitting was performed on the STA; for movies, we fitted the parameters of a Gaussian using gradient descent, initializing with a mean based on the STA location and a standard deviation of 1. We added one more inductive bias on top of the regularization techniques. We (4) modified the final default softplus nonlinearity by adding learnable parameters to it: *parameterized nonlinearity (PNL)* (Equation 1), allowing the model to learn a more nuanced relationship between linear filter output and spiking response.

On all datasets, we drastically improved performance by adding combinations of the inductive biases (1)–(4) to the models (Fig. 2C). The best resulting model on salamander noise data incorporated all inductive biases and achieved a median correlation of 0.41 (IQR 0.25–0.52), which is 0.32 points better than the unregularized model. For the marmoset data, the best setting differed for the two stimuli. The best model for the white noise stimulus included all the biases and improved from the baseline median correlation of 0.16 to 0.59 (IQR 0.31–0.69). For the movie stimulus, the best model included all biases but the Gaussian fitting. It also showed a substantial improvement – from a baseline median correlation of 0.4 to 0.73 (IQR 0.62–0.83). Thus, the LN models are decent predictive models for both noise and movie stimuli when properly regularized.

We also examined how each single bias (1)–(4) improved the performance of LN models. To achieve this, we calculated the difference in performance between two models: one with and one without applying a given bias, while all other biases were the same between the two models. We found that all biases improved performance across species and stimulus-types at least in certain combinations (Fig. 2D–F). For salamander noise, spatial cropping, space-time separation, and Gaussian fitting were all effective regularization methods. Marmoset noise models mainly benefited from space-time separation, Gaussian fitting, and spatial cropping. Marmoset movie models primarily benefited from a parameterized nonlinearity. The Gaussian fitting was not beneficial in this case; rather, it lowered performance.

The effect of specific methods varied depending on the presence of other inductive biases. For the white noise stimulus, the critical factor was reducing the number of parameters – the specific method of parameter reduction was not as decisive (Fig. 2G). For natural movies, parameter reduction also had a positive influence. However, the key factor was fitting the nonlinearity parameters.

### The effect of an inductive bias greatly varies across datasets and stimuli

In addition to examining overall average effects, we also analyzed how different inductive biases influence performance on a per-dataset basis.

For the salamander data, the results were consistent across datasets. All inductive biases had a generally positive effect, and performance improved steadily as more biases were applied (Tab. 1). The marmoset data revealed a more nuanced pattern. While white noise performance generally improved with additional inductive biases for Datasets 1 and 2, we observed a substantial performance drop in Dataset 4 when including the Gaussian fitting regularization (Tab. 2, last two rows). Aside from this notable exception, the trend of improved performance with more biases held for the marmoset white noise stimuli.

**Table 1.** LN model for white noise salamander. Mean LN model performance on white noise salamander data with different combinations of inductive biases. SC stands for spatial-cropping, S-T for space-time separation, PNL for parameterized nonlinearity, and GF for Gaussian fitting. X means inductive bias was not applied, ✓means it was applied. Columns R1 – R5 refer to the five different datasets.

| LN model version |  |  |  | Mean over cells | R1 | R2 | R3 | R4 | R5 |
| --- | --- | --- | --- | --- | --- | --- | --- | --- | --- |
| SC | S-T | PNL | GF |  |  |  |  |  |  |
| X | X | X | X | 0.1 | 0.17 | 0.08 | 0.07 | 0.06 | 0.08 |
| ✓ | X | X | X | 0.22 | 0.3 | 0.2 | 0.19 | 0.18 | 0.18 |
| X | ✓ | X | X | 0.2 | 0.32 | 0.19 | 0.16 | 0.14 | 0.15 |
| X | ✓ | ✓ | X | 0.23 | 0.36 | 0.2 | 0.16 | 0.12 | 0.19 |
| ✓ | ✓ | X | X | 0.3 | 0.4 | 0.3 | 0.26 | 0.25 | 0.25 |
| ✓ | ✓ | ✓ | X | 0.36 | 0.47 | 0.36 | 0.29 | 0.25 | 0.3 |
| X | ✓ | ✓ | ✓ | **0.38** | 0.48 | 0.37 | 0.32 | 0.3 | 0.34 |
| ✓ | ✓ | ✓ | ✓ | **0.38** | 0.48 | 0.36 | 0.32 | 0.3 | 0.33 |

**Table 2.** LN model for white noise marmoset. Mean LN model performance on white noise marmoset data with different combinations of inductive biases. SC stands for spatial-cropping, S-T for space-time separation, PNL for parameterized nonlinearity, and GF for Gaussian fitting. X means inductive bias was not applied, ✓means it was applied.

| LN model version |  |  |  |  | Mean over cells | Dataset 1 | Dataset 2 | Dataset 4 |
| --- | --- | --- | --- | --- | --- | --- | --- | --- |
|  | SC | S-T | NL | GF |  |  |  |  |
| 1. | X | X | X | X | 0.2 | 0.14 | 0.15 | 0.43 |
| 2. | ✓ | X | X | X | 0.32 | 0.25 | 0.27 | 0.59 |
| 3. | X | ✓ | X | X | 0.37 | 0.29 | 0.28 | 0.66 |
| 4. | X | ✓ | ✓ | X | 0.44 | 0.4 | 0.27 | 0.69 |
| 5. | ✓ | ✓ | X | X | 0.46 | 0.44 | 0.28 | 0.68 |
| 6. | ✓ | ✓ | ✓ | X | 0.49 | 0.44 | 0.39 | 0.72 |
| 7. | X | ✓ | ✓ | ✓ | 0.5 | 0.59 | 0.43 | 0.3 |
| 8. | ✓ | ✓ | ✓ | ✓ | **0.55** | 0.59 | 0.43 | 0.55 |

The results on marmoset natural movies were less uniform (Tab. 3). Fitting a Gaussian degraded performance on Datasets 1 and 4, but improved it on Datasets 2 and 3 – more so on Dataset 3 – (rows 3&4 vs. rows 7&8). The impact of space-time separation was also dataset-specific: it improved performance on Datasets 1 and 3, but had a detrimental effect on Datasets 2 and 4 (row 1 vs. row 3). Interestingly, the effect of a given bias also depended on the type of stimulus. For example, space-time separation improved predictions on white noise for Datasets 2 and 4, yet reduced performance on natural movies for the same datasets. The pattern for Gaussian fitting was even less consistent. For Dataset 1, Gaussian fitting was beneficial on white noise, but it hurt performance on natural movies. In contrast, for Dataset 4, Gaussian fitting consistently reduced performance across both stimulus types, and for Dataset 2, it improved performance across both stimulus types.

**Table 3.** LN model for natural movies marmoset. Mean LN model performance on natural movies marmoset data with different combinations of inductive biases. SC stands for spatial-cropping, S-T for space-time separation, PNL for parameterized nonlinearity, and GF for Gaussian fitting. X means inductive bias was not applied, ✓means it was applied.

| LN model version |  |  |  |  | Mean over cells | Dataset 1 | Dataset 2 | Dataset 3 | Dataset 4 |
| --- | --- | --- | --- | --- | --- | --- | --- | --- | --- |
|  | SC | S-T | PNL | GF |  |  |  |  |  |
| 1. | X | X | X | X | 0.41 | 0.41 | 0.26 | 0.38 | 0.59 |
| 2. | ✓ | X | X | X | 0.44 | 0.34 | 0.38 | 0.53 | 0.7 |
| 3. | X | ✓ | X | X | 0.43 | 0.53 | 0.17 | 0.39 | 0.43 |
| 4. | ✓ | ✓ | X | X | 0.51 | 0.6 | 0.25 | 0.45 | 0.51 |
| 5. | X | ✓ | ✓ | X | 0.56 | 0.53 | 0.3 | 0.61 | 0.8 |
| 6. | ✓ | ✓ | ✓ | X | **0.64** | 0.6 | 0.5 | 0.67 | 0.85 |
| 7. | X | ✓ | X | ✓ | 0.4 | 0.41 | 0.34 | 0.56 | 0.24 |
| 8. | ✓ | ✓ | X | ✓ | 0.4 | 0.41 | 0.37 | 0.56 | 0.25 |
| 9. | X | ✓ | ✓ | ✓ | 0.55 | 0.51 | 0.54 | 0.7 | 0.49 |
| 10. | ✓ | ✓ | ✓ | ✓ | 0.57 | 0.54 | 0.5 | 0.72 | 0.57 |

Taken together, these results indicate that there is no single inductive bias or combination that yields optimal performance across datasets – even within the same species and stimulus condition. This stresses the importance of tuning the LN model design to the specific characteristics of each dataset.

### Nonlinear CNNs improve predictive performance

Based on the results in the previous sections, we saw that reducing the number of learnable parameters boosts performance. While we explored a range of regularization techniques on the level of single-cell models, another possibility is to use a different model design that allows us to model a population of neurons simultaneously. In such a case, a portion of the parameters does not have to be learned individually for each neuron; instead, they can be shared by all neurons. To achieve this, we used CNN models with a *core-readout* architecture (Fig. 3A). Such models have been used successfully in V1 as well as for RGCs cell prediction (McIntosh et al., 2016; Klindt et al., 2017; Lurz et al., 2021; Höfling et al., 2024). They consist of one or more layers of convolutions – the *core* – which creates a feature space shared by all neurons. The *readout* then allows each neuron to learn its receptive field position in this feature space and weight the shared features at this position (Fig. 3A).

**Figure 3.**
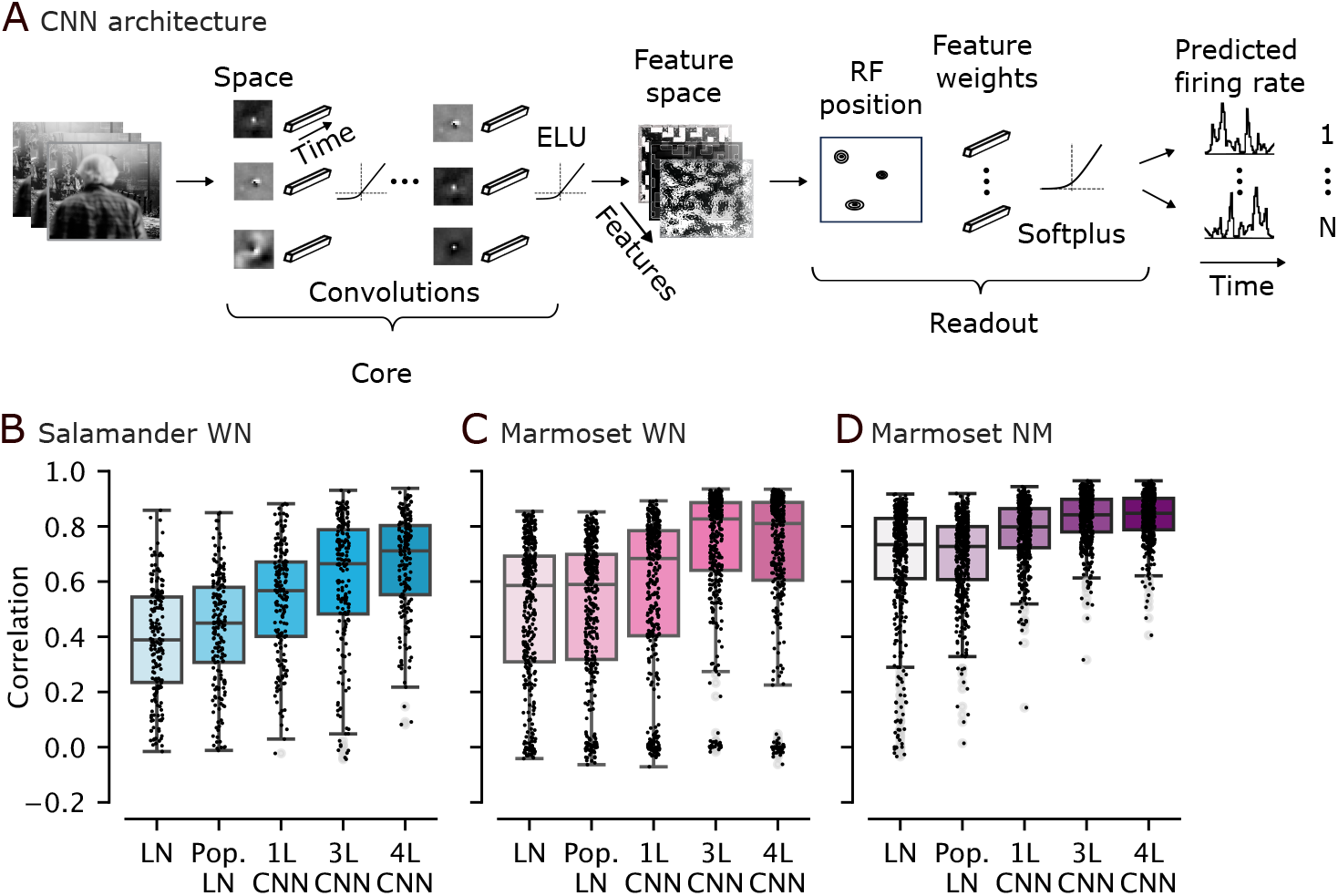
**A**. The core-readout architecture of the population LN and CNN models. *N* equals the number of neurons. **B.-D**. The IQ1-IQ3 range (boxplots) with median (black line) of model types of increasing complexity on **B**. salamander cells being shown noise stimuli, **C**. marmoset cells being shown noise stimuli, **D**. marmoset cells being shown movie stimuli. Small black dots represent cells.

We first built an LN model in this fashion, which we refer to as a population LN model. It has only one space-time separated convolutional layer in the core, directly followed by the readout. The only nonlinearity is at the very end after the readout. Since there are no nonlinearities between the spatial and temporal convolutions and between the core and the readout, this model is effectively an LN model as well, with the restriction that the linear filter is constructed as a linear combination of a basis set shared by all the modeled cells.

The sizes of the spatial and temporal convolutional filters were, along with all other hyperparameters, established by evaluating performance on the validation set. The best spatial filter size values were larger compared to the cropped versions of single cell LN models (Tab. 4–Tab. 6).

**Table 4.** Best model hyperparameters for models trained on salamander RGC responses to noise.

| **Salamander on WN** | Spat. input kernel | Temp. input kernel | Spat. hidden kernel | Temp. hidden kernel | Spat. dilations | Channels |
| --- | --- | --- | --- | --- | --- | --- |
| Population LN | 17 | 25 | - | - | 1 | 16 |
| CNN 1 | 17 | 25 | - | - | 1 | 64 |
| CNN 3 | 11 | 25 | 5 | 5 | 1, 2, 2 | 16, 32, 64 |
| CNN 4 | 11 | 25 | 9 | 5 | 1, 1, 1, 1 | 8, 16, 32, 64 |

**Table 5.** Best model hyperparameters for models trained on marmoset RGC responses to noise.

| **Marmoset on WN** | Spat. input kernel | Temp. input kernel | Spat. hidden kernel | Temp. hidden kernel | Spat. dilations | Channels |
| --- | --- | --- | --- | --- | --- | --- |
| Population LN | 27 | 35 | - | - | 1 | 32 |
| CNN 1 | 27 | 35 | - | - | 1 | 32 |
| CNN 3 | 21 | 35 | 3 | 5 | 1, 1, 1 | 16, 32, 64 |
| CNN 4 | 25 | 25 | 3 | 5 | 1, 1, 1, 1 | 8, 16, 32, 64 |

**Table 6.** Best model hyperparameters for models trained on marmoset RGC responses to movies.

| Marmoset on NM | Spat. input kernel | Temp. input kernel | Spat. hidden kernel | Temp. hidden kernel | Spat. dilations | Channels |
| --- | --- | --- | --- | --- | --- | --- |
| Population LN | 27 | 35 | - | - | 1 | 32 |
| CNN 1 | 25 | 35 | - | - | 1 | 32 |
| CNN 3 | 21 | 35 | 5 | 3 | 1, 1, 1 | 16, 32, 64 |
| CNN 4 | 17 | 27 | 3 | 5 | 1, 1, 2, 3 | 8, 16, 32, 64 |

We trained this model end-to-end for all three data types. We found that the population LN out-performed the best regularized single-cell LN model on the salamander, improving from a median correlation of 0.39 (IQR 0.23–0.54) to 0.45 (IQR 0.31–0.58). On the marmoset data, the performance of the single cell and population LN was comparable: 0.59 (IQR 0.31–0.69) and 0.59 (IQR 0.32–0.7) for the single cell and population LN respectively on marmoset noise and 0.73 (IQR 0.61–0.83) and 0.73 (IQR 0.61–0.8) on marmoset movies for single cell and population LN respectively (Fig. 3B–D).

While the predictive performance of the single-cell and population LN models was good, a substantial gap remains in predicting the RGCs’ responses perfectly. So far, we have tried to improve performance by regularizing linear models. However, numerous studies show that the retina performs nonlinear computations (Baccus et al., 2008; Gollisch and Meister, 2010; Kastner and Baccus, 2014; Kuo et al., 2016; Turner and Rieke, 2016; Karamanlis and Gollisch, 2021). To push performance further, we moved beyond LN models and implemented models with multiple nonlinearities in successive processing layers, allowing them to learn these nonlinear computations. The core-readout CNN model structure allowed us to easily incorporate nonlinear transformations, placing them after each space-time convolutional layer within the core and between the core and the readout. We implemented networks of various depths to establish the depth that yields the highest performance and establish benchmarks at multiple complexity levels. Our CNNs ranged from one to four layers.

The complexity of the CNN architecture, particularly the number of layers and subsequent nonlin-earities, directly impacted performance (Fig. 3B–D). Models with a single nonlinearity after the core have already demonstrated a significant improvement over the population LN model, indicating the value of nonlinear processing in these response prediction tasks and, thus, in retinal processing. As the number of layers – and therefore also the number of nonlinear transformations – increased, we observed a corresponding increase in performance. For salamander data, performance continued to increase up to four layers, reaching a median correlation of 0.71 (IQR 0.55–0.8) across cells. For the marmoset data, performance saturated at three layers, with a median correlation of 0.83 (IQR 0.64–0.88) across cells for noise and 0.84 (IQR 0.78–0.9) across cells for movies. This trend suggests that the depth of the network, which relates to its ability to form more complex representations and feature hierarchies, is a critical factor in achieving accurate predictions of ganglion cell responses. Notably, the absolute performance gain between linear and nonlinear models on natural movies is smaller than on white noise.

### All models fail to fully generalize across stimulus statistics

We have established that LN models and, more so, CNNs can predict RGC responses to both noise and movies with high accuracy, mimicking retinal computations under the given stimulus statistics. We next asked whether these computations are specifically adapted to the stimulus statistics or whether these models generalize to other stimuli. To do so, we tested how well models trained on one stimulus type (noise or movies) predict the responses of RGCs to the other stimulus (movies or noise). If the models maintain their predictive performance across both stimulus types, it would suggest that the retinal computations are universal, without specific adaptation to the type of stimulus. If prediction accuracy is maintained in only one direction, it would imply that the given training stimulus encompasses a more universal set of features, capturing aspects of both stimulus types. If model generalization is unsuccessful in both directions, it would indicate a tailored adaptation of the receptive fields to the unique statistics of each stimulus.

For this analysis, we used only marmoset data where we recorded the same neurons under both stimulus conditions. We trained all models included in this analysis on the same number of stimulus frames for both noise and movie data.

We took models trained on one of the stimuli and evaluated them on the other in two scenarios. The first scenario we evaluated was unadapted prediction on the other stimulus, that is, we evaluated the model directly on the type of stimulus it had not seen during training, without adapting any of the model’s weights. Under these conditions, neither stimulus ensemble allowed models to reach in-domain performance on the other stimulus type (Fig. 4A, B; white vs. colored bars). This was true across all models, but the gap was larger for the CNN models than the LN models. Notably, the unadapted CNN’s performance in this scenario even dropped below that of the unadapted LN model. These results suggest that the processing of neurons adapts to the stimulus statistics, and the adaptation is strongest for nonlinear response properties, as they do not generalize across varying stimulus statistics.

**Figure 4.**
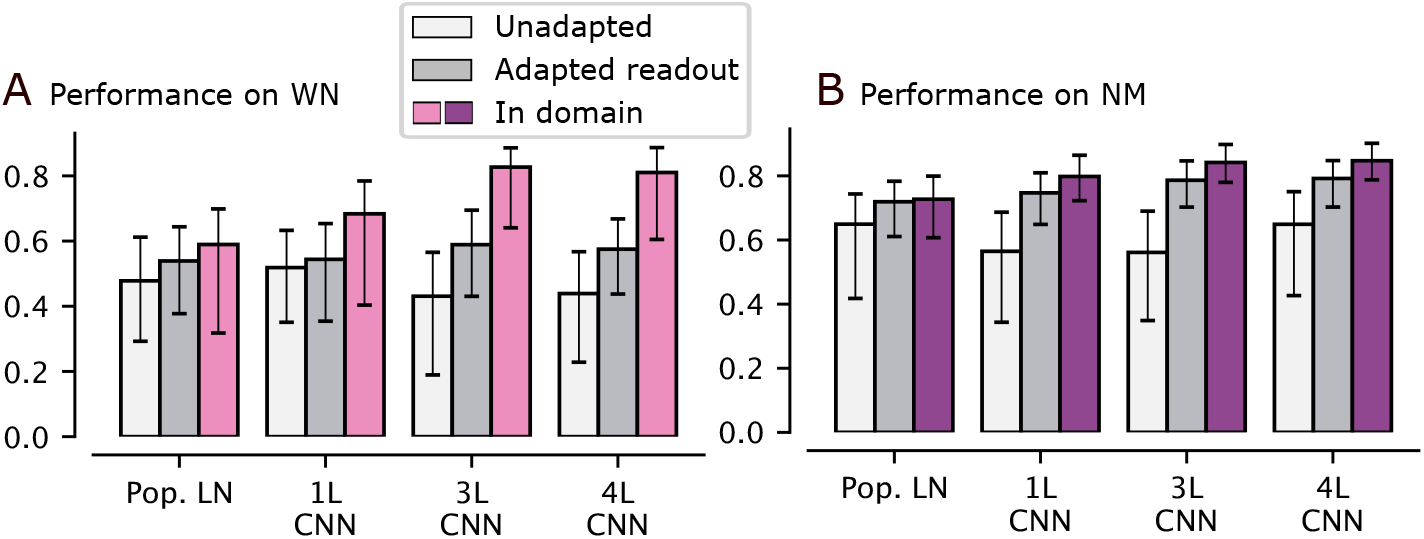
**A**. Models evaluated on white noise. White: Undadapted performance – i.e., models trained on natural movies applied directly to white noise data. Gray: performance with adapted readout: model trained on natural move, readout, and parameterized final nonlinearity adapted by training on noise data. Pink: In domain: Model trained on white noise evaluated on white noise. **B**. Models evaluated on natural movies. White: Unadapted performance - i.e., model trained on white noise applied directly to natural movies. Gray: performance with adapted readout: model trained on white noise, readout, and parameterized final nonlinearity adapted by training on natural movie data. Purple: In domain: Model trained on natural movies evaluated on natural movies.

Next, we asked whether the features learned by the shared core generalized across stimulus statistics, or whether generalization fails because of simple adaptation phenomena that can be accounted for by linearly recombining the learned features. To test this idea, we allowed the models to adapt their linear readout. We kept the weights of the core fixed and fine-tuned the weights of the readout and the output nonlinearity on the other stimulus. Adapting the readout somewhat improved generalization from movies to noise, but a fairly large gap remained (Fig. 4A; gray vs. colored bars). In contrast, the model trained on noise generalized reasonably well to movies when allowing the readout to adapt (Fig. 4B; gray vs. colored bars), almost closing the gap for the LN model. This result suggests that while the linear filters change across stimulus statistics, the basis set of features learned in the core from white noise spans most of the space learned to predict movie data. In contrast, the features from the movie data do not span a vector space that can fully account for white noise responses.

## Discussion

We benchmarked a range of carefully optimized models, ranging from LN to CNN models, for predicting the responses of RGCs across two species – marmosets and salamanders and two stimulus ensembles – white noise and natural movies.

We explicitly chose to compare LN models and CNNs because they represent two distinct ends of the complexity spectrum in computational modeling of retinal encoding. LN models are simple, highly interpretable, and have historically been widely adopted by neuroscientists due to their transparency and computational efficiency (Chichilnisky, 2001; Rust et al., 2005). As such, they serve as classical baseline models. In contrast, more recent CNNs have demonstrated superior predictive performance in various neural response prediction tasks (McIntosh et al., 2016; Klindt et al., 2017; Lurz et al., 2021; Sinz et al., 2018), largely benefiting from their flexible architectures that learn to capture complex nonlinearities in a data-driven manner, albeit at the cost of reduced interpretability. We acknowledge that several intermediate modeling strategies exist, employing various inductive biases and structural constraints, which blend simplicity and predictive power (Pillow et al., 2008; McFarland et al., 2013; Freeman et al., 2015; Maheswaranathan et al., 2018). However, evaluating the models at the two ends of the spectrum – LN as the simplest and CNNs as the most complex – provides a clear framework for benchmarking intermediate models, facilitating an intuitive assessment of their relative contributions and effectiveness. In the future, we plan to integrate these models into the newly emerging collaborative initiative openretina (D’Agostino et al., 2025), which aims to streamline model use and comparison, making benchmarking even more accessible.

### The tuning of linear-nonlinear models

Our results demonstrate that the performance of LN models varies substantially depending on their construction. This raises concerns about their widespread use as baselines (McIntosh et al., 2016; McFarland et al., 2013; Freeman et al., 2015; Sridhar et al., 2025; Gogliettino et al., 2024). In our experiments, incorporating standard inductive biases into an LN model often led to performance improvements of over twofold on the same dataset. Given that these biases are not universally beneficial across datasets or stimulus types, comparisons between LN baselines and more complex models may be misleading if the LN model is not regularized and tuned appropriately. This reinforces the need for rigorous hyperparameter optimization when using LN models as benchmarks and highlights the importance of standardized evaluations on publicly available datasets to ensure fair comparisons.

The heterogeneity in the effect of inductive biases across datasets likely reflects biological variability – such as differences in the composition of recorded cell types. Among the biases tested, two were particularly variable in their effects: space-time separability and Gaussian spatial filtering. Both affect the structure of the receptive field surround, which may explain their inconsistent utility. In the case of space-time separation, our implementation imposed a rank-one constraint on the linear filter, effectively forcing the center and surround to share a common temporal profile. However, it is known that the center typically lags slightly behind the surround (Kilavik et al., 2003; Cowan et al., 2016). This mismatch may lead to degraded performance in datasets where the role of the surround is more pronounced. Similarly, Gaussian fitting imposes a strong prior by modeling the spatial receptive field with a single Gaussian. This constraint effectively suppresses the surround altogether. This may explain why Gaussian fitting negatively impacts the performance of more datasets on natural movie stimuli, where surround contributions are more prominent than in white noise conditions (Vystrčilová et al., 2025).

### Natural vs. white noise stimuli

Training models on natural movie stimuli is vital for studying visual processing, as white noise inputs have been shown to fail to drive key nonlinear retinal computations such as contrast adaptation, motion extrapolation, and omitted stimulus responses (Tanaka et al., 2019; Maheswaranathan et al., 2023). While some previous work suggested that LN models are unable to capture RGC responses under natural stimulation (Heitman et al., 2016), Vystrčilová et al. (2025) demonstrated that LN models can actually predict responses to natural stimuli fairly well. Moreover, they found that LN models generalize better from white noise to natural movies than in the reverse direction. We confirm this finding and further show that the same asymmetry holds for more complex CNN models. The directionality of generalization we observe is in contrast with findings in primary visual cortex, where generalization from natural movies to white noise was found to be better than the reverse (Sinz et al., 2018).

The asymmetry may reflect differences in stimulus structure: natural movies contain less high-frequency spatial and temporal content, which makes it harder for models trained on them to learn the more high-resolution features needed to predict responses to white noise. This could explain the reduced generalization from movies to noise, despite the likely richer computations being elicited by naturalistic input.

Such findings seem to challenge the rationale for using natural movie stimuli. However, models trained on white noise may generalize better in terms of predictive performance while still failing to engage key retinal computations. Generalization accuracy does not directly indicate whether the underlying computational mechanisms are captured.

Generally, metrics based on explained variance are somewhat tricky to interpret. While they reach unity if and only if one has a perfect model, differences in explained variance do not necessarily tell us how interesting this explained variance is. The nonlinear features of retinal processing (Schwartz et al., 2007, 2012; Kastner and Baccus, 2014; Maheswaranathan et al., 2023) are significant; however, since they do not occur frequently in natural scenes, accurately accounting for them will provide only minor improvements in terms of explained variance.

In our data, responses to movies are predicted more accurately than responses to white noise, likely due to the more substantial luminance fluctuations in the movie stimuli, which account for a greater portion of the variance in natural movie stimuli. This is not always the case, McIntosh et al. (2016) report closer absolute prediction performance values with slightly better results on white noise for both LN models and CNNs. Thus, absolute differences in predictive performance may not fully reflect differences in underlying neural computation.

Lastly, although natural-movie training has revealed interesting nonlinear response properties absent in noise-trained models (Maheswaranathan et al., 2023), the reverse might also hold: we may simply not have identified properties that are not learned from movies. Thus, neither stimulus is inherently “better”. As of now, it appears that retinal processing flexibly adapts to the statistics of the input provided, and we still lack a constructive, low-parametric, or even parameter-free model of this adaptation.

## Methods

### Data

#### Salamander data

We used five datasets from adult axolotl salamander retinas, each comprising the responses of RGCs to spatiotemporal white noise stimuli, measured using multi-electrode arrays (MEAs). The stimuli were structured into trials, with each trial consisting of a 50-second non-repeating training segment followed by a 10-second test segment that was repeated across trials. A total of 39 to 248 trials were recorded per dataset. The acquisition of these five datasets has been previously described in Liu et al. (2017), and they are publicly available (Gollisch and Liu, 2018).

#### Marmoset data

We also used four datasets recorded using MEAs from the retinas of adult common marmosets (*Callithrix Jacchus*) from Sridhar et al. (2025). Each dataset consisted of responses of RGCs under spatiotemporal white noise and natural movie stimulation with jittering, which mimicked the statistics of natural eye saccades and fixations. For natural movie stimulation, we used an adapted version of a short, publicly available science-fiction film by the Blender Foundation under the CCA 3.0 license (Tears of Steel, accessible at https://mango.blender.org/). Each trial of the white noise stimulus consisted of 150 seconds of non-repeating frames and 30 seconds of repeating frames, whereas each trial of the natural movie stimulus consisted of 300 seconds of repeating frames and 60 seconds of non-repeating frames. The stimuli were shown at a frame rate of 85 Hz. Three datasets contained responses to the same 10 trials of the white noise stimulus, and one dataset contained different 10 trials of the white noise stimulus. *Dataset 1, Dataset 3* and *Dataset 4* contained 20 trials of the natural movie stimulus, whereas *Dataset 2* contained 10 trials. The acquisition and preprocessing of all datasets have been described in Sridhar et al. (2025). In brief, the retina was extracted from the eye and mounted on an MEA while under constant perfusion of an oxygenated solution, ensuring minimal light contamination of the tissue. An oLED screen focused on the photoreceptor layer of the mounted tissue was used to project the stimulus while the activity of RGCs was recorded using the MEA setup. Spiking responses of individual RGCs were isolated using a semi-automated spike-sorting pipeline and binned at the resolution of the stimulus refresh rate for further modeling.

#### Cell selection

We only used cells with reliable responses to the stimuli for our analyses. The reliability of the cells was determined based on the fraction of explainable variance (Heitman et al., 2016; Cadena et al., 2019) in each cell’s responses to the repeating segments of the stimuli across trials. All cells had to exceed a threshold of 0.15 to be used for further analysis. For the marmoset datasets, we retained all cells that exceeded this threshold for both white noise and natural movie stimuli. This left us with 186 reliable cells from the salamander, specifically 62, 53, 23, 29 and 19 cells for the five different retinas, and 384 reliable cells across *Dataset 1* (235 cells), *Dataset 2* (69 cells) and *Dataset 4* (80 cells) of the marmoset retinas. All cells in *Dataset 3* were deemed unreliable on the white noise stimulus. Therefore, we used 91 cells that responded reliably to the natural movie stimulus and used this dataset only for experiments involving natural movies.

### Linear-nonlinear models

#### White noise models

Filters for each cell for the white noise model were estimated using spike-triggered averaging (STA). We calculated the STA as

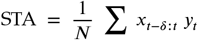

where *x*_*t*_ is the frame presented at time *t, y*_*t*_ is the spike-count recorded at time-point *t* and *N* is the recorded number of spikes. *δ* - the size of the temporal dimension of the STA was 25 for salamander and 30 for marmoset. To this baseline filter, we then apply the following inductive biases:

- *Spatial-crop* around receptive field – The original STA has size 60×60 pixels spatially for salamander and 40×40 pixels spatially for marmoset. These sizes can capture the complete receptive field both spatially and temporally. However, such large crops require many parameters to be fit, which can lead to overfitting. Therefore, we cropped more tightly around the receptive field to reduce the number of parameters. Specifically, we used crop sizes of 20×20 pixels for salamander and 15×15 pixels for marmoset.
- *Space-time separation* – To get a space-time separation, we calculate the STA as above. We then computed the singular value decomposition and used the first resulting decomposition kernels as the spatial and temporal filters.
- *Gaussian fitting* – Another way to reduce the number of parameters and prevent over-fitting is fitting a Gaussian to the spatial filter of the LN model. We estimated the spatial filter as described in space-time separation and then fitted a 2D Gaussian to it. The values that were more than two standard deviations away from the mean were set to zero. The temporal filter remained the same as in the space-time separated version.
- *parameterized nonlinearity* – As the default output nonlinearity, we used the softplus function. This can be parameterized by adding learnable parameters *α* and *β* as follows:

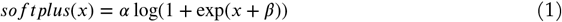

where *x* is the input. We optimized these parameters using Poisson loss (Equation 2) between predicted firing rate *p* and recorded firing rate *r* using stochastic gradient descent.

#### Natural movie models

Natural movie stimulus has spatial and temporal correlations. We, therefore, estimated the entire LN model filter learning it end-to-end, minimizing the Poisson loss (Equation 2) between recorded firing rate *r* and the predicted firing rate *p* using stochastic gradient descent (Vystrčilová et al., 2025). We applied the same biases as for the white noise model filters with the following differences:

- *Spatial-crop* – The original STA had size 40×40 pixels spatially. We cropped tightly around the receptive field. Specifically, we used 15×15 pixels, the same as the crops for the marmoset noise models.
- *Gaussian fitting* – We randomly initialized the temporal filter and the mean and covariance matrix of a non-isotropic Gaussian. The Gaussian then parameterized the spatial component of the filter. We trained all the learnable parameters using stochastic gradient descent.

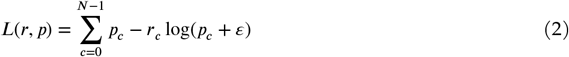

#### Training

For all parameters optimized with stochastic gradient descent in LN models, we used the following training scheme, similar to Vystrčilová et al. (2025). We split the trials in our datasets into 80% train and 20% validation splits. We randomly initialized the learnable parameters and updated them by minimizing the Poisson loss (Equation 2) between predicted *p* and recorded *r* response values. We trained on the training trials for up to 1,000 epochs, with early stopping triggered when validation correlation has not increased for 30 epochs. The initial learning rate was 3*e*^−3^, and we employed the Pytorch `ReduceLROnPlateau` learning rate scheduler with a patience of 15 on the validation correlation with a minimal learning rate of 1*e*^−7^. The optimizer we used was `Adam` (Kingma and Ba, 2017).

#### Evaluation

We report the final performances of the best models based on validation correlation by calculating the correlation coefficient for a given RGC *c* (Equation 3) between the predictions *p*_*c*_ and the trial averaged firing rates ⟨*r*_*c*_⟩ on the held-out test sequence with length *T*.

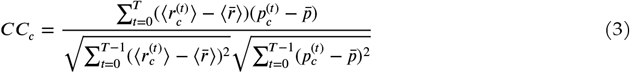

### CNN models

#### Architecture

We trained multiple models of various depths. All of them had a similar core-readout structure as in Höfling et al. (2024). The core consisted of a varying number of layers of space-time separated convolutions. After each spatio-temporal convolution, we applied an Exponential Linear Unit (ELU) nonlinearity (Clevert et al., 2015) and a batch normalization (Ioffe and Szegedy, 2015). We used different optimal hyperparameters for each combination of CNN depth, species, and stimulus type. The best hyperparameter values are based on validation performance in Tab. 4 – Tab. 6. The kernels of the first layer were smoothed by applying a 2D (3×3 pixels) Laplace filter in the spatial dimensions and a 1D (1×3 pixels) Laplace filter in the temporal dimension. In contrast to (Höfling et al., 2024), temporal kernels were not parameterized as Fourier series but optimized directly. In the readout, each cell’s receptive field was modeled as an isotropic Gaussian from which we sample at training time and take the mean at inference time. The response function was modeled as an affine function of the core’s weighted feature maps at the receptive field positions, followed by a parameterized softplus (Equation 1). The vectors used to weight the features for the modeled neurons were regularized using the L1 Norm.

The architectural feature that seemed most significant for good model performance was making the first layer spatial and temporal kernels larger relative to those in subsequent layers. Presumably, the large kernel allows the models to capture the entire receptive field, including the surround. Regularization, on the other hand, had a minor effect on model performance.

#### Training

We split the trials in the datasets in the same way as for the LN models, into 80% train and 20% validation splits. We trained the CNNs by minimizing the Poisson loss (Equation 2) on the training trials for up to 1000 epochs, with early stopping when the correlation on the validation trials had not increased for 50 consecutive steps. The initial learning rates were 0.008 for natural movies and 0.006 for white noise. We employed the ReduceLROnPlateau learning rate scheduler from PyTorch, with a patience of 15 on the validation correlation and a minimal learning rate of 1*e*^−6^. The optimizer was Adam (Kingma and Ba, 2017).

#### Evaluation

For each complexity level, i.e., number of layers, we selected the best model based on the estimated validation correlation and report its final performances by calculating the correlation coefficient (Equation 3) between the model predictions *p* and the trial averaged responses ⟨*r*⟩ of the held-out test sequence of the dataset.

#### Cross-stimulus evaluation

To evaluate cross-stimulus generalization, we used the marmoset datasets *Dataset 1, Dataset 2*, and *Dataset 4*, which contain responses to both white noise and natural movie stimuli of the same cells. We used only five trials of natural movie data from these datasets when training models for the cross-stimulus generalization analysis to match the size of the white noise dataset, because the natural movie trials were twice the length of the white noise trials. In the unadapted setting, models trained on one stimulus type were directly applied to the other to assess the correlation between predicted and recorded spike counts. In the adapted readout setting, the readout weights and final nonlinearity parameters of models trained on one stimulus were adjusted by training on the other stimulus, following the same procedure as for the initial training.

## Acknowledgements

MFB thanks the International Max Planck Research School for Intelligent Systems (IMPRS-IS).

## Author Contributions

Conceptualization: M.V., A.E., T.G; Data Curation: S.S, M.V, M.K, D.K, H.S, V.R, S.K, S.Z, M.M, Formal Analysis: M.V, S.S; Funding Acquisition: A.E, T.G; Investigation: M.V., S.S; Methodology: M.V., S.S. M.B.; Resources: S.S, M.K, D.K, H.S, V.R, S.K, S.Z; Software: M.V., S.S; Supervision: A.E., T.G.; Visualization: M.V.; Writing - Original Draft Preparation: M.V., A.E., M.B; Writing - Review & Editing: M.V., S.S., A.E., M.B., T.G.

## Footnotes

1 https://neuralprediction.org/npc/index.php

